# Clusterin in Alzheimer’s disease: an amyloidogenic inhibitor of amyloid formation

**DOI:** 10.1101/2021.04.02.438207

**Authors:** Panagiotis M. Spatharas, Georgia I. Nasi, Paraskevi L. Tsiolaki, Marilena K. Theodoropoulou, Nikos C. Papandreou, Andreas Hoenger, Ioannis P. Trougakos, Vassiliki A. Iconomidou

**Affiliations:** Section of Cell Biology and Biophysics, Department of Biology, School of Sciences, National and Kapodistrian University of Athens, Panepistimiopolis, Athens 157 01, Greece; University of Colorado at Boulder, Department of Molecular, Cellular and Developmental Biology, Boulder, CO 80309-0347, USA

**Keywords:** Clusterin, Molecular chaperones, Aggregation-prone regions, Amyloid, Amyloid-*β*, Amyloid inhibitors, Alzheimer’s disease

## Abstract

Clusterin is a heterodimeric glycoprotein (α- and β-chain), which has been described as an extracellular molecular chaperone. In humans, clusterin is an amyloid associated protein, co-localizing with fibrillar deposits in several amyloidoses, including Alzheimer’s disease. To clarify its potential implication in amyloid formation, we located aggregation-prone regions within the sequence of clusterin α-chain, via computational methods. We had peptide-analogues of each region chemically synthesized and experimentally demonstrated that all of them can form amyloid-like fibrils. We also provide evidence that the same peptide-analogues can inhibit amyloid-β fibril formation. These findings elucidate parts of the molecular mechanism in which clusterin inhibits amyloid formation. At the same time, they hint that molecular chaperones with amyloidogenic properties might have a role in the regulation of amyloid formation, essentially acting as functional amyloids.

## Introduction

Since the 17th century, physicians had already started to describe clinical disorders that were later included in a broad group of diseases, nowadays known as amyloidoses (Kyle, 2011). Prime examples are Alzheimer’s disease (AD), Parkinson’s disease, type 2 diabetes and AL amyloidosis (Benson et al, 2020). It is widely considered that these diseases are closely associated with the accumulation and close packing of normally soluble proteins, which end up creating highly ordered, insoluble aggregates. The so called amyloid fibrils, are deposited extracellularly in organs or tissues, causing significant damage (Benson et al., 2020).

Molecular chaperones can inhibit amyloid formation in its early stages, thus preventing amyloid-related cytotoxicity (Wentink et al, 2019). Recent studies though, have shown that molecular chaperones can have amyloidogenic properties (Ecroyd & Carver, 2009; Ecroyd et al, 2010; Niewold et al, 1999; Tsiolaki et al, 2017), despite their seemingly contradicting physiological function. *In vitro* experiments performed by our team have hinted that clusterin could be such a protein (Tsiolaki et al., 2017). Clusterin is a ubiquitous, conserved mammalian glycoprotein, whose main isoform is secreted and has been described as an extracellular molecular chaperone (Jones & Jomary, 2002). Human clusterin precursor is a 449 residue-long polypeptide chain, which, after having its signal peptide removed, is reduced to 427 residues, and goes through several post-translational modifications (Jones & Jomary, 2002). Mature clusterin is cleaved in two chains —*α*-chain, consisting of 222 residues and *β*-chain, consisting of 205 residues— with five disulfide bonds forming between them (Jones & Jomary, 2002). Its molecular weight is approximately 80 kDa, almost 30% of it being carbohydrates, added by glycosylation (Jones & Jomary, 2002). Clusterin’s structure has yet to be experimentally determined, but is believed to contain three long molten globule-like regions and five amphipathic *α*-helices, which allow for hydrophobic interactions with its client-proteins (Bailey et al, 2001).

Under normal circumstances, clusterin exists in solution as heterogeneous aggregates (*Poon et* al, 2002). It has been suggested that, at mildly acidic pH (Poon et al., 2002), these aggregates disassociate and the disassociated chaperone-active subunits function in an ATP-independent manner (Poon et al, 2000). They bind misfolded client-proteins and form a high molecular weight complex. The formation of the complex allows for stabilization of the misfolded proteins, which are consequently refolded by other molecular chaperones (Bailey et al., 2001) (visualized in Fig 1). Clusterin’s ever-growing list of known clients includes a variety of proteins, such as cellular receptors, apolipoproteins, the complement complex, immunoglobulins, amyloid-forming proteins and non-protein molecules, like lipids and heparin (Calero et al, 2005).

**Figure 1.**
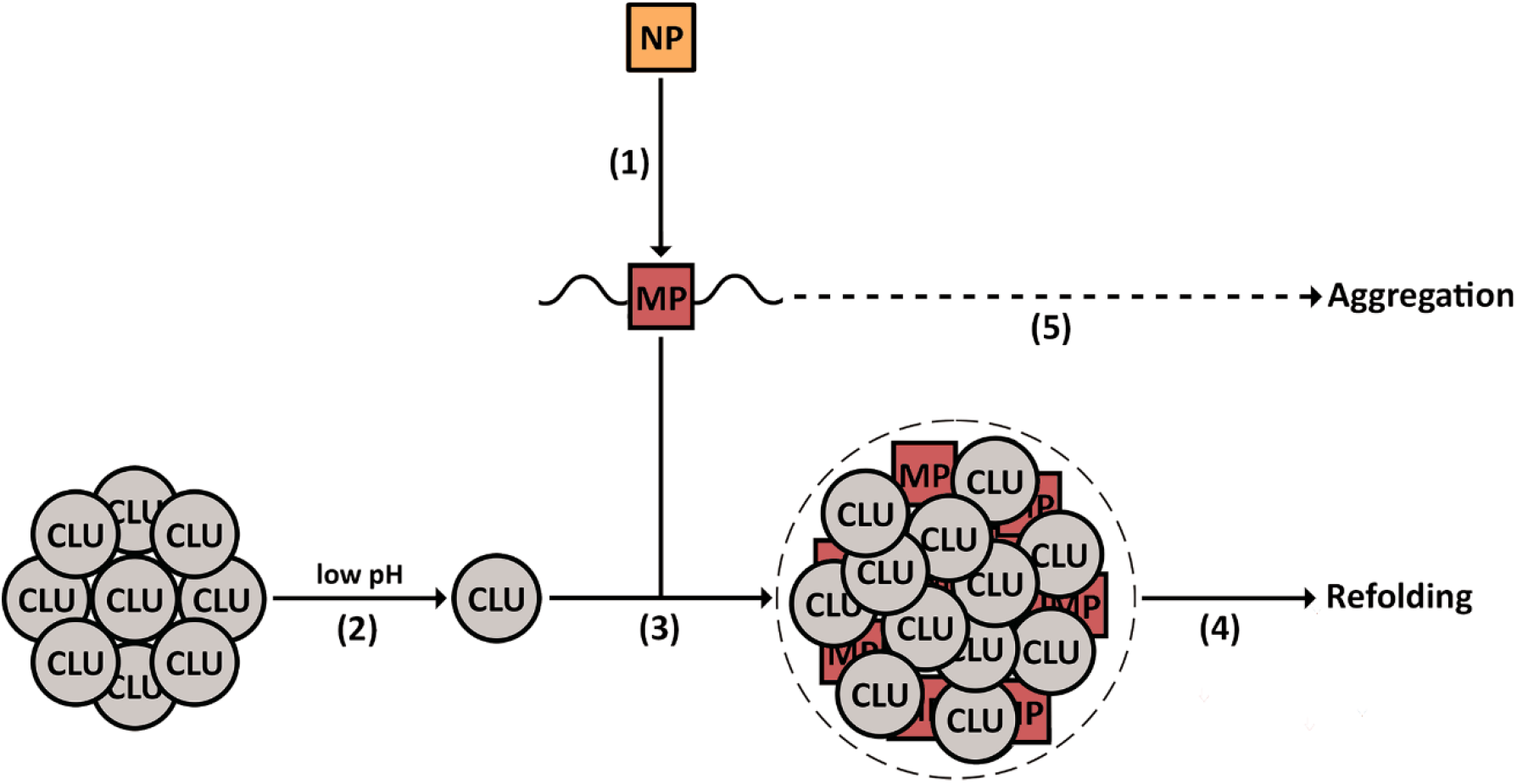
Representation of clusterin’s molecular chaperone activity. (1) Under stress (low pH), native proteins (NP) may partially unfold and become misfolded. (2) At slightly acidic pH, clusterin’s (CLU) oligomers start disassociating and the chaperone-active clusterin subunits get released. (3) The free clusterin subunits bind the misfolded proteins (MP) and form a complex, effectively stabilizing them, (4) while other molecular chaperones refold them to their native state. (5) If this mechanism proves ineffective, the misfolded proteins may form aggregates.

Having such a broad range of clients, it is only logical that clusterin contributes in many physiological and pathological processes (Rohne et al, 2016). AD is one of the most notable pathologies in which clusterin is involved (Calero et al., 2005). Consistent with its chaperone function, clusterin keeps the amyloid-*β* (A*β*) peptide soluble while transporting it in biological fluids, modulates its permeation through the blood-brain barrier and contributes to its clearance, effectively halting amyloid formation (Matsubara et al, 1996). In direct contradiction to that, it has been reported that clusterin contributes to the early stages of AD pathogenesis (DeMattos et al, 2002; Oh et al, 2019). To this day, clusterin’s role in AD is yet to be fully understood. Despite the mystery surrounding clusterin’s implication in AD, it is a known fact that it co-localizes with A*β* fibrillar deposits (Calero et al., 2005). This is also true for other amyloidoses, in which clusterin is an amyloid associated protein (Nastou et al, 2019).

It can be assumed that clusterin is found in the amyloid deposits because it is drifted along by its client-proteins. In this study, we examined a different scenario, one that doesn’t rule out the possibility of clusterin being amyloidogenic itself. We used AMYLPRED (Frousios et al, 2009), a consensus algorithm for the prediction of amyloid propensity, in order to identify aggregation-prone regions in clusterin *α*-chain. We had peptide-analogues of each region chemically synthesized and experimentally demonstrated that all of them can form amyloid-like fibrils *in vitro*. The same peptide-analogues, despite being amyloidogenic, can inhibit A*β* fibril formation. Based on our findings, we proposed a putative mechanism in which clusterin prevents amyloid formation. The suggested mechanism could also explain the contradictory reports that hint at clusterin’s implication in the acceleration of the appearance of AD symptoms. At the same time, we hope that the basis of clusterin’s amyloid inhibiting activity could give insight into the implementation of peptide-based amyloidogenesis inhibitors in the treatment of amyloidoses.

## Results

### All wild-type clusterin *α*-chain peptide-analogues form amyloid-like fibrils *in vitro*

According to the prediction of AMYLPRED, clusterin *α*-chain has five aggregation-prone regions. The peptide-analogues that correspond to those regions are NFHAMFQ, AMDIHF, ILSVD, YYLRVTT and EVVVKLF, as depicted in Fig 2. NFHAMFQ has previously been studied by our lab and proved to exhibit amyloidogenic properties *in vitro* (Tsiolaki et al., 2017). Our current findings reveal that AMDIHF, ILSVD, YYLRVTT, and EVVVKLF can also be characterized as amyloidogenic *in vitro*, making a total of five experimentally verified aggregation-prone regions in clusterin *α*-chain. Characterization was based on the tinctorial criteria, which are commonly used for the identification of amyloid fibrils (transmission electron microscopy, X-ray diffraction from protein fibers, ATR FT-IR spectroscopy and Congo Red birefringence assay) (Sunde & Blake, 1998).

**Figure 2.**
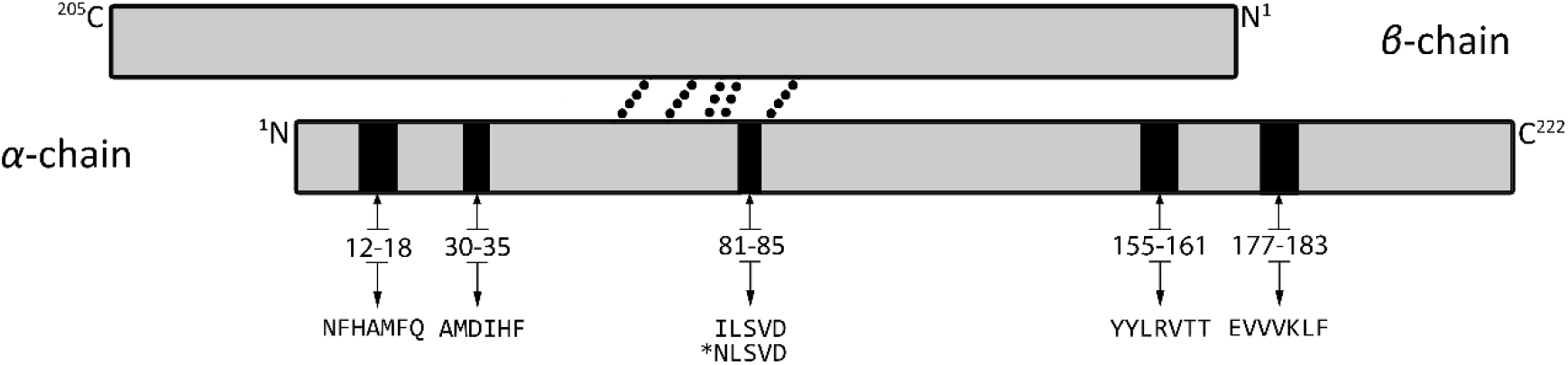
A simple representation of clusterin’s two chains. Each chain is colored grey, with a black outline, and its sequence is numbered as found in the mature protein. The *α*-chain consists of 222 amino acid residues. The *β*-chain is slightly shorter, consisting of 205 amino acid residues. The two chains are joined by five disulfide bonds, depicted as black dots. The aggregation-prone regions of the *α*-chain are depicted as black rectangles on clusterin’s sequence.

Transmission electron micrographs show that AMDIHF, ILSVD, YYLRVTT, and EVVVKLF self-assemble into amyloid-like fibrils (Fig 3 A-D, respectively). AMDIHF fibrils exhibit considerable polymorphism, forming both supercoils and tapes (Fig EV1 A-C), while EVVVKLF fibrils form large twisting ribbons, with a 162 nm pitch (Fig EV1D). ILSVD forms amyloid-like fibrils that are packed more densely in comparison to the other peptides (Fig 3), indicating that it could be the most aggregation-prone region in clusterin α-chain.

**Figure 3.**
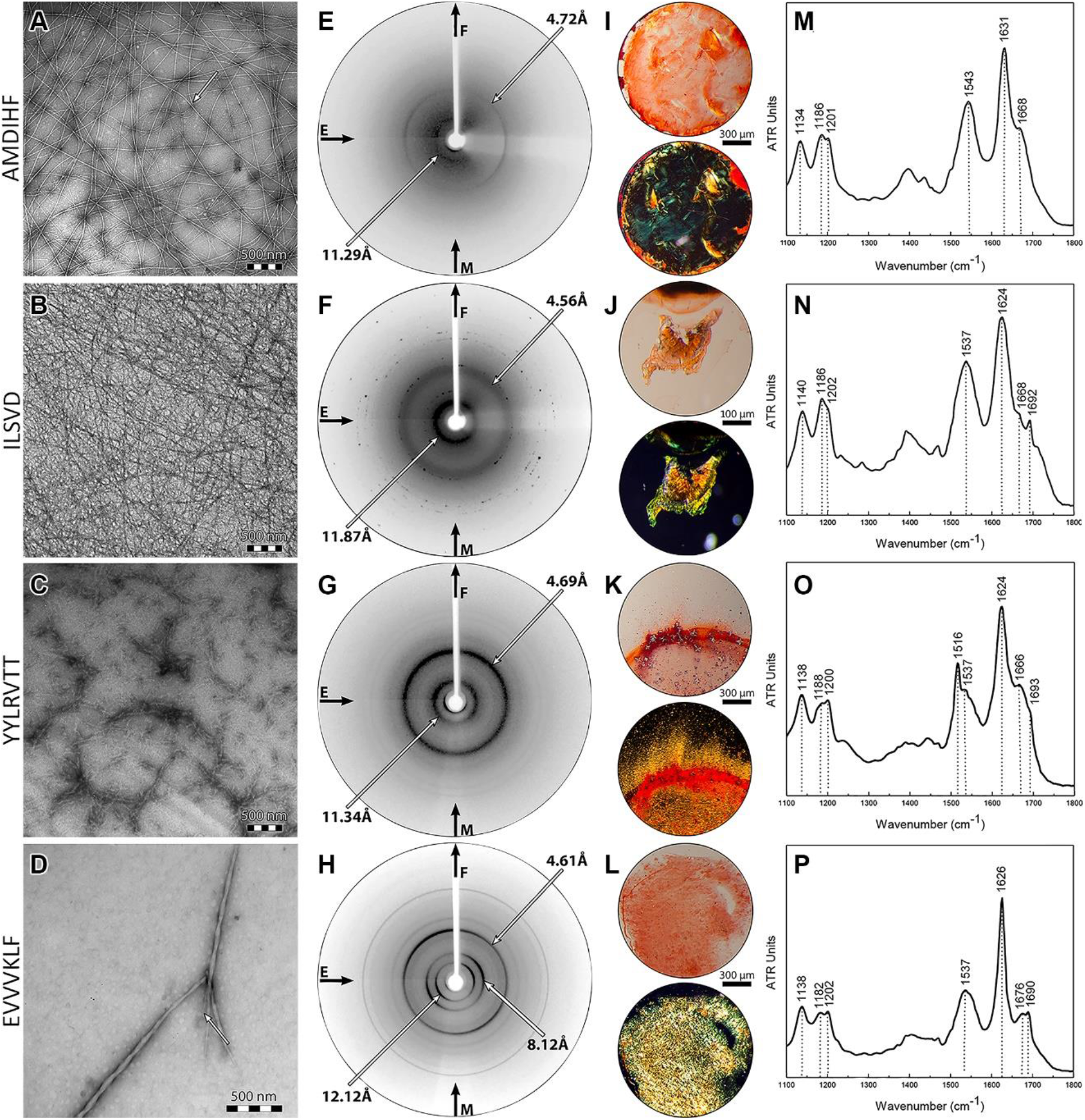
All the peptide-analogues fulfill the experimental criteria. (A-D) Transmission electron micrographs of amyloid-like fibrils derived from 500 μm solutions of peptides AMDIHF, ILSVD, YYLRVTT and EVVVKLF, respectively. White arrows mark single fibrils with the diameter of typical amyloid fibrils. (E-H) X-ray diffraction patterns of oriented fibers, derived from 1 mM solutions of the four peptides, respectively. Reflections marked with arrows are indicative of the “cross-*β*” structure, which amyloids typically have. (I-L) Photomicrographs of gels, derived from 500 μM solutions of the four peptides, respectively. Congo red is bound, as seen under bright field illumination (upper). The apple-green birefringence that amyloids typically exhibit is observed under crossed polars (lower). (M-P) ATR FT-IR spectra (1100–1800 cm^-1^) produced from thin hydrated films created by 500 μM solutions of the four peptides, respectively. Fibrils derived from each peptide has *β*-sheet secondary structure, as hinted by the presence of strong amide I and II bands.

Oriented fibers formed by each peptide solution, produce X-ray diffraction patterns, indicative of the presence of the “cross-*β*” structure. Most reflections are ring-like, most likely because the fibrils are not perfectly aligned in a parallel manner (Makin & Serpell, 2005). The exact measurements for each of the four diffraction patterns can be seen in Fig 3 (E-H). It is worth pointing out that EVVVKLF diffraction pattern has two equatorial reflections, one at 8.12 and one at 12.12 Å. Upon closer examination of the twisting ribbons, formed by EVVVKLF (Fig EV1D), the maximum width is observed in the middle of the pitch, and measures approximately 53 nm, while the minimum width is observed at the site of the twist, and measures approximately 35 nm. The quotient of the two equatorial reflections (8.12: 12.12 ≈ 0.67) is roughly equal to the quotient of the minimum and maximum ribbon width (35: 53 ≈ 0.66). The equatorial reflections are indicative of the distance between packed beta sheets. Thus, EVVVKLF fibrillar structures are probably formed by β-sheets that are more tightly packed at the twist, in comparison to the middle of the ribbon. The periodicity of the tightening and untightening of the β-sheets is revealed by a combination of X-ray and TEM data.

The dominance of the *β*-sheet secondary structure is supported by ATR FT-IR spectroscopy data, since the spectra of all four peptide-analogues reveal bands that can be assigned to *β*-sheets (Surewicz et al, 1993) (Table 1 and Fig 3 M-O).

**Table 1.**
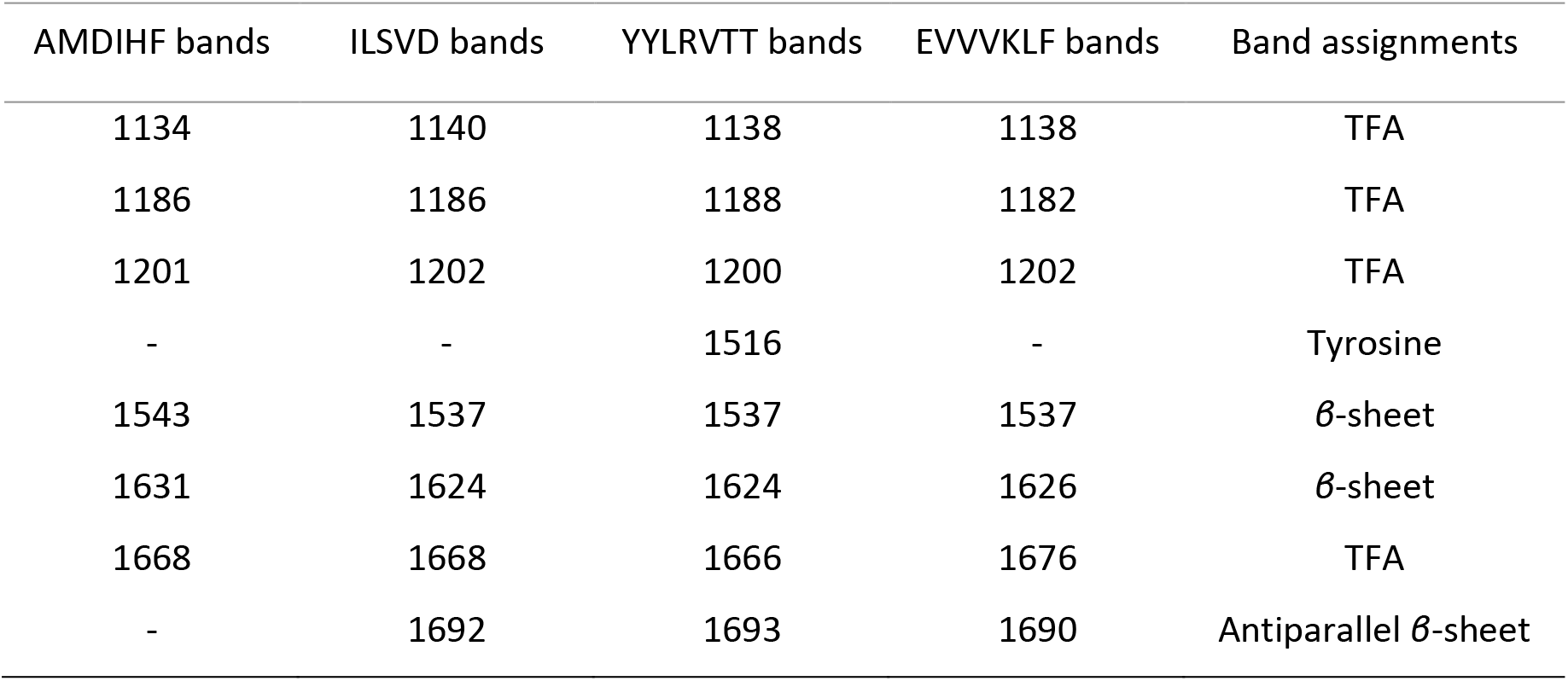
Bands observed in the ATR FT-IR spectra obtained from thin hydrated films produced by AMDIHF, EVVVKLF, YYLRVTT and ILSVD solutions, respectively, and their tentative assignments.

Every peptide binds Congo Red, as seen under bright field illumination in a polarizing stereomicroscope (Fig 3 I-L, upper). When the polars are crossed, the peptides exhibit the apple-green birefringence (Fig 3 I-L, lower) that amyloids typically exhibit.

### NLSVD, a mutant counterpart of ILSVD, does not exhibit amyloidogenic properties

Recent studies have identified a novel pathological mutation on clusterin’s gene, in AD patients. In particular, p.I360N changes the aggregation-prone region ^81^ILSVD^85^ to ^81^NLSVD^85^ (Bettens *et al*, 2015). The resulting sequence is not predicted by AMYLPRED, hinting that it is not aggregation-prone. To experimentally confirm that the mutated region is not aggregation-prone, we had NLSVD chemically synthesized. Indeed, transmission electron microscopy (TEM) shows that NLSVD does not self-assemble into amyloid fibrils (Fig EV2A). Also, NLSVD does not form oriented fibers, suitable for X-ray diffraction, hinting that there are no fibrils in its solution. Finally, when stained with Congo Red and stereoscopically observed between crossed polars, it does not exhibit apple-green birefringence (Fig EV2B).

### Clusterin *α*-chain peptide-analogues inhibit *Aβ*_42_ amyloid formation *in vitro*

Full-length clusterin is known to inhibit A*β* fibril formation *in vitro* (Yerbury et al, 2007). We aimed to pinpoint segments of its sequence that could act as catalysts of said inhibition. Each clusterin peptide-analogue was co-incubated with equimolar amounts of A*β*_42_ and the aggregation kinetics were evaluated with Thioflavin T (ThT) fluorescence measurements over time and TEM.

ThT fluorescent measurements reveal that individual A*β*_42_ starts forming fibrils almost immediately and reaches maximal fluorescence signal after approximately 7 hours. When co-incubated with clusterin peptide-analogues (A*β*_42_+NFHAMFQ, A*β*_42_+AMDIHF, A*β*_42_+ILSVD, A*β*_42_+YYLRVTT, and A*β*_42_+EVVVKLF), A*β*_42_ exhibits decreased emission. ILSVD and especially EVVVKLF seem to be the most potent inhibitors (Fig 4J and 4P, respectively). As previously shown, individual ILSVD is incredibly amyloidogenic, as is A*β*_42_. The fact that interaction between the two leads to decreased fibril formation, hints that peptides with high amyloidogenic potential, could negate each other’s fibril-forming properties. EVVVKLF bears some sequence similarity with ^16^KLVFFA^21^, a key amyloid-forming segments of A*β*_42_ (Lu et al, 2019). Considering that peptide-analogues derived from ^16^KLVFFA^21^ have been suggested as potential therapeutic agents for the treatment of AD (Chafekar et al, 2007), it is not surprising that EVVVKLF has the highest inhibiting effect among the five peptides. NFHAMFQ and AMDIHF also seem to inhibit A*β*_42_ fibril formation (Fig 4F and 4H, respectively), but their potential as inhibitors is less significant than that of ILSVD and EVVVKLF. YYLRVTT effectively delays fibril formation, but after approximately 20 hours, fluorescence signal begins to rise (Fig 4N). After 40 hours, A*β*_42_+YYLRVTT reaches fluorescence intensity levels which are comparable to those of individual A*β*_42_.

**Figure 4.**
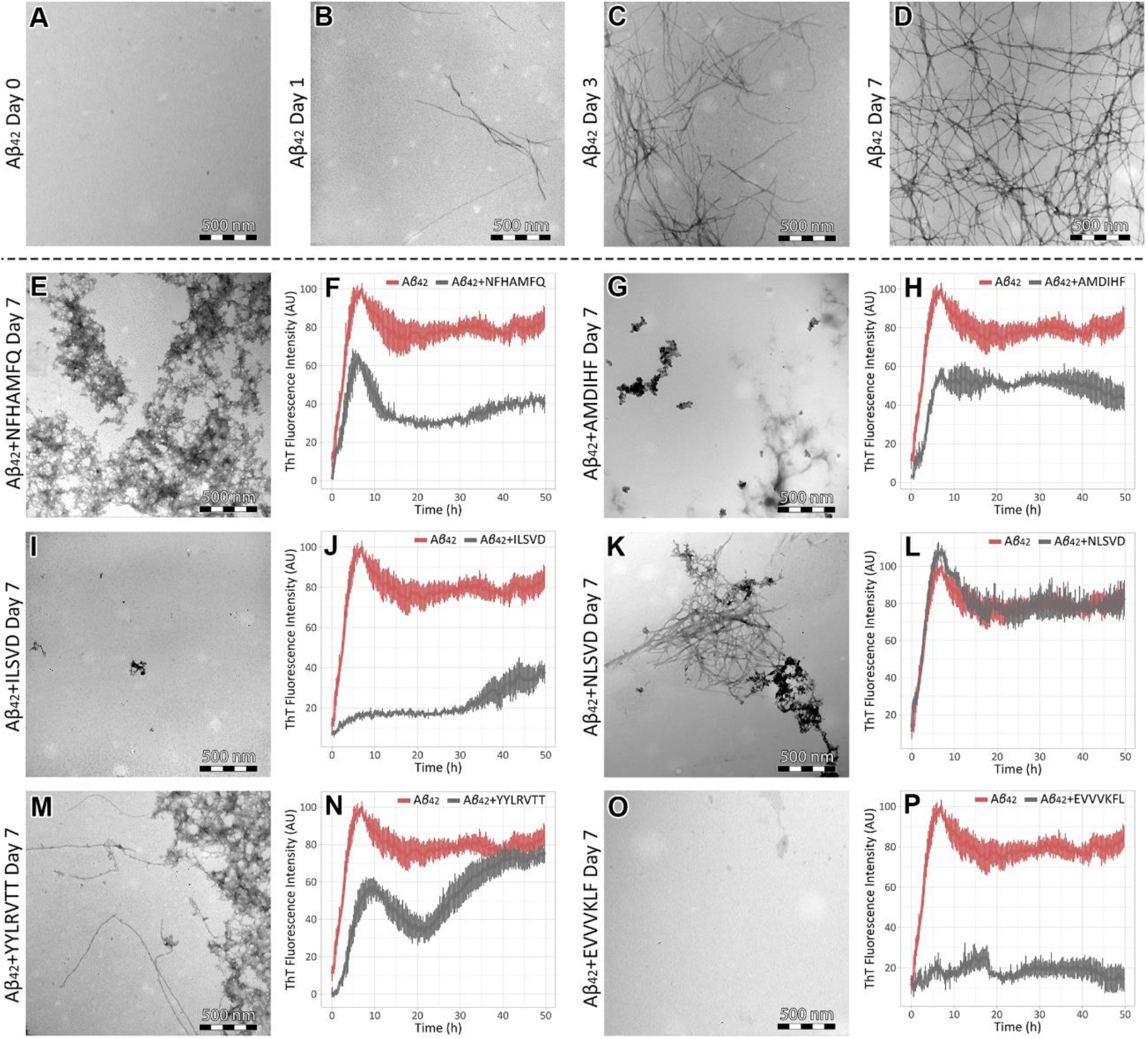
Clusterin peptide-analogues inhibit or delay A*β*_42_ fibril formation. (A-D) Transmission electron micrographs of A*β*_42_, incubated for 0, 1, 3 and 7 days, respectively. Amyloid fibrils are observed from day 1 to day 7. The number of fibrils raises as incubation time increases. (E, F) Transmission electron micrograph and ThT fluorescence emission spectrum of A*β*_42_ co-incubated with NFHAMFQ for 7 days and over a period of 50 hours, respectively. (G, H) Transmission electron micrograph and ThT fluorescence emission spectrum of A*β*_42_ co-incubated with AMDIHF for 7 days and over a period of 50 hours, respectively. (I, J) Transmission electron micrograph and ThT fluorescence emission spectrum of A*β*_42_ co-incubated with ILSVD for 7 days and over a period of 50 hours, respectively. (K, L) Transmission electron micrograph and ThT fluorescence emission spectrum of A*β*_42_ co-incubated with the mutant peptide, NLSVD, for 7 days and over a period of 50 hours, respectively. (M, N) Transmission electron micrograph and ThT fluorescence emission spectrum of A*β*_42_ co-incubated with YYLRVTT for 7 days and over a period of 50 hours, respectively. (O, P) Transmission electron micrograph and ThT fluorescence emission spectrum of A*β*_42_ co-incubated with EVVVKLF for 7 days and over a period of 50 hours, respectively. Error bars in ThT fluorescence emission spectra represent standard deviation among triplicates. TEM experiments contained 100 μM A*β*_42_ and 100 μM clusterin peptide-analogues, while ThT experiments contained 10 μM and 10 μM, respectively.

TEM confirms that all five peptide-analogues inhibit or delay A*β*_42_ fibril formation, after 7 days of incubation. When individually incubated, A*β*_42_ forms straight and unbranched fibrils with indefinite length and a diameter of approximately 80 Å (Fig 4 B-D). A small number of fibrils is observed after 1 day of incubation (Fig 4B). The population of fibrils increases significantly after 3 days (Fig 4C) and reaches its maximum after 7 days (Fig 4D). As expected, ILSVD and EVVVKLF prove to be the most potent inhibitors (Fig 4I and 4O, respectively). The entire grids were thoroughly examined, and no fibrils were observed. NFHAMFQ and AMDIHF also inhibit A*β*_42_ fibril formation, as fibrils were not observed anywhere on the grid (Fig 4E and 4G, respectively). However, both seem to induce the formation of amorphous aggregates. YYLRVTT seems to delay, but not inhibit fibril formation. In contrast to the other peptides, fibrils can be observed (Fig 4M). The number of fibrils is similar to that of individual A*β*_42_, after 1 day of incubation. Much like NFHAMFQ and AMDIHF, YYLRVTT also seems to induce the formation of amorphous aggregates.

On the contrary, ThT kinetics and TEM show that NLSVD, the mutant counterpart of ILSVD, does not inhibit A*β*_42_ amyloid formation *in vitro*. A*β*_42_ and A*β*_42_+NLSVD fluorescence curves mostly overlap, while A*β*_42_+NLSVD show a slightly enhanced fluorescence peak (Fig 4K). Transmission electron micrographs reveal that A*β*_42_ fibril morphology is different to that of A*β*_42_+NLSVD. A*β*_42_+NLSVD fibrils, after 7 days of incubation, are more densely packed and slightly wider than those of individual A*β*_42_, after the same incubation period (Fig 4L and Fig EV2C, white arrow). Furthermore, Aβ42+NLSVD fibrils tend to interact laterally and form loosely bound tapes (Fig EV2C, blue arrows).

### Computational insight into the inhibition of A*β*_42_ fibril formation by EVVVKLF

To shed light on the interaction between A*β*_42_ and EVVVKLF, the most potent of the five peptide-inhibitors, molecular dynamics simulations were performed, using an NMR structure of pentameric A*β*_42_ (PDB ID: 2BEG) (Luhrs et al, 2005). A structure for EVVVKLF was generated through the “Builder” tool in PyMOL (Delano, 2005). Molecular docking was performed through the automated protein docking server ClusPro (Kozakov et al, 2013; Kozakov et al, 2017; Porter et al, 2017; Vajda et al, 2017). The generated clusters were evaluated with the balanced scoring scheme. Following energy minimization, the interaction between A*β*_42_ key amyloid-forming segment ^17^LVFFA^21^ (Fig 5A, dark grey) and EVVVKLF (Fig 5A, light grey) was ascertained (Fig 5A, left). A*β*_42_+EVVVKLF complex structural stability was monitored through per-residue root mean square fluctuation (RMSF) calculation, as well as time-dependent root mean square deviation (RMSD) measurements (Fig 5B and 5C, respectively). After 600 ns of molecular dynamics simulations, a complex dissociation has occurred, significantly changing the conformation of the pentamer, in comparison to its original state (Fig 5A). The C-termini of all five chains are characterized by large fluctuations (0.4 – 0.8 nm), with chains-A and -B exhibiting the highest mobility. In contrast, the N-termini of the aforementioned chains exhibit the lowest mobility among all chains, while chain-E N-terminus is characterized by fluctuations of over 0.8 nm. Secondary structure analysis was performed using the dictionary of secondary structure of proteins (DSSP) algorithm (Kabsch & Sander, 1983; Touw et al, 2015) and reveals that *β*-sheet content gradually decreases (Fig 5D). Intermolecular hydrogen bonds are formed, even before the simulation has begun, and reach a maximum number of 12 (Fig 5E). Upon examination of the last frame of the simulation (Fig 5A, right), EVVVKLF has partly adopted a *β*-strand conformation. Using UCSF Chimera and the “FindHBond” tool (Pettersen et al, 2004), we show that EVVVKLF forms three possible hydrogen bonds with the chain-A ^16^LVFFA^21^ region, becoming a part of an intermolecular β-sheet. EVVVKLF also forms hydrogen bonds with the C-terminus of the A-chain. By forming hydrogen bonds with both regions, it essentially stiches the N-terminal ^17^LVFFA^21^ to the C-terminus. The fact that RMSF values are low for the N-terminus and high for the C-terminus hints that the later moves towards the former. In this way, fibril elongation from chain-A could be blocked. The addition of more than one peptide-inhibitors could promote interactions with chain-E, effectively blocking both elongation epitopes of the pentamer.

**Figure 5.**
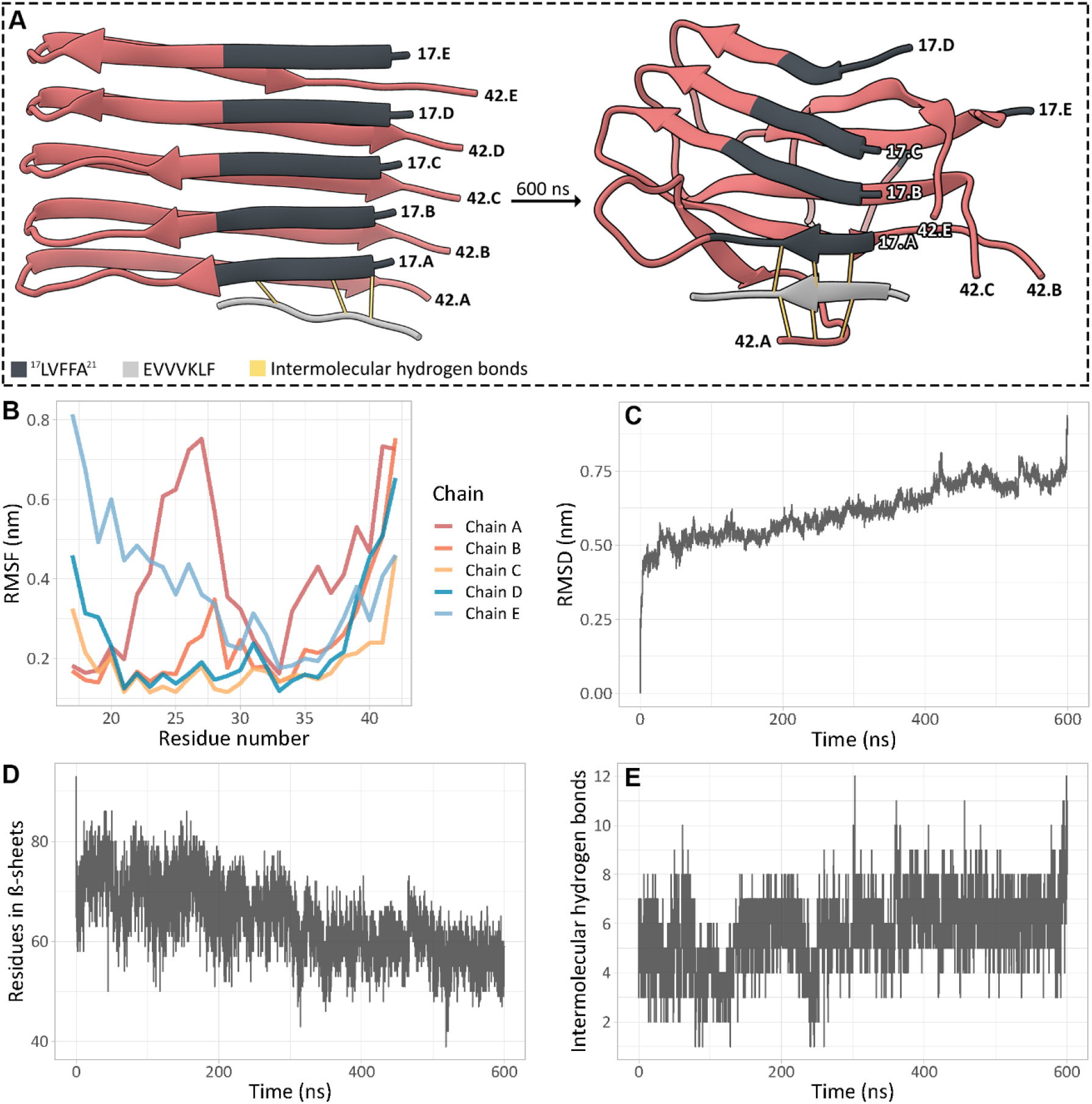
Results of molecular dynamics simulations for A*β*_42_+EVVVKLF. (A) First (0 ns, left) and last (600 ns, right) frames of the molecular dynamics simulation. The position of the N- and C-terminals of each A*β*_42_ chain are marked on each frame (17.A-17.E and 42.A-42.E, respectively). EVVVKLF (light grey) forms hydrogen bonds (yellow lines) with the chain-A ^17^LVFFA^21^ (dark grey), right after molecular docking and energy minimization. More bonds are formed by the end of the simulation. (B) RMSF per residue plot, after 600 ns of molecular dynamics simulations. (C) RMSD over time plot. (D) Total number of residues in *β*-sheets, according to DSSP, over time. (E) Number of intermolecular hydrogens bonds, between A*β*_42_ and EVVVKLF, over time.

### Natively disordered regions in human clusterin α-chain

Human clusterin is believed to have three long molten globule-like regions (Bailey et al., 2001). PONDR^®^ VLTX (http://www.pondr.com) predicts five natively disordered regions in clusterin α-chain (Fig 6), the longest of which is located at the C-terminus. All five aggregation prone regions are located near predicted natively disordered regions. ^12^NFHAMFQ^18^ is on the right of the N-terminal natively disordered region and on a putative amphipathic *α*-helix (*Bailey et al*., 2001). Putative *α*-helices are believed to be implicated in the interaction between clusterin and its client proteins (Bailey et al., 2001). ^30^AMDIHF^35^ is located directly to the right of said putative amphipathic *α*-helix. ^81^ILSVD^85^ is surrounded by two natively disordered regions. ^155^YYLRVTT^161^ is also located directly to the left of a natively disordered region, while ^177^EVVVKLF^183^ is partly predicted as natively disordered. Interestingly, the EVVVKLF peptide-analogue adopts *β*-sheet secondary structure, according to the ATR FT-IR spectrum (Fig 3P). The proximity of all five aggregation-prone regions to natively disordered regions may hint that, due to the flexibility of the later, the former could act semi-independently of the protein.

**Figure 6.**
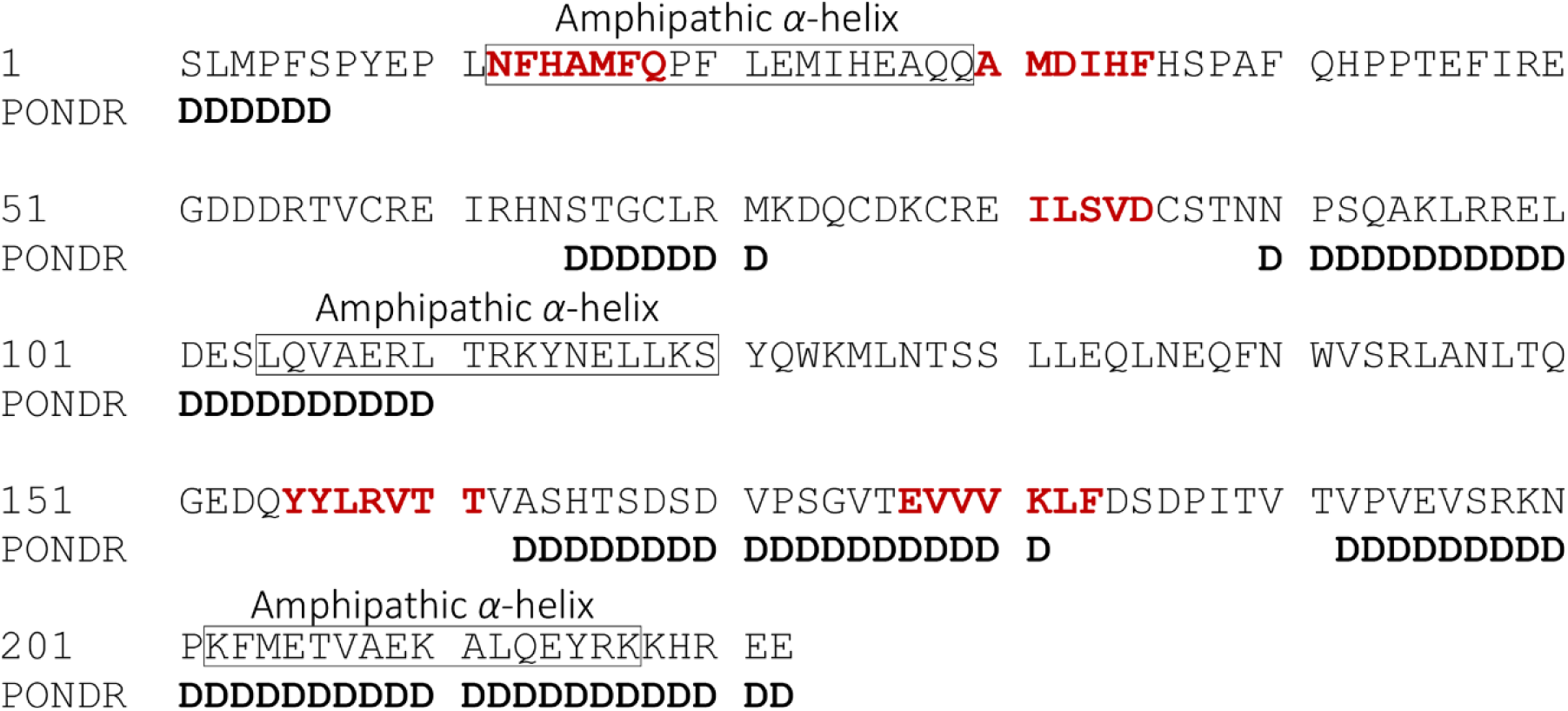
Relative position of natively disordered regions (as predicted by PONDR^®^ VLTX), aggregation-prone regions, and putative amphipathic *α*-helices, on the sequence of human clusterin *α*-chain. All aggregation-prone regions are located near predicted natively disordered regions. ^12^NFHAMFQ^18^ is located on a putative amphipathic *α*-helix, while ^177^EVVVKLF^183^ is partly predicted as natively disordered.

## Discussion

### Full-length clusterin could also have amyloidogenic properties *in vitro*

Recent studies have shown that short aggregation-prone segments, known as amyloidogenic determinants, can lead to amyloid formation, even when they are inserted in non-amyloidogenic proteins, via protein engineering (Teng & Eisenberg, 2009). The fact that clusterin has five such segments in its *α*-chain, provides potent indication that the *α*-chain and consequently the full-length protein, may intrinsically exhibit amyloidogenicity. This argument has fascinating implications, especially considering clusterin’s role as a molecular chaperone.

### Molecular chaperones and their involvement in amyloid formation and inhibition

One of the fundamental functions of molecular chaperones is considered to be the prevention of protein aggregation and consequently, amyloid fibril formation (Wentink et al., 2019). Based on that notion, it has been suggested that molecular chaperones could be used as anti-amyloid drugs (Yerbury & Kumita, 2010). In direct contradiction to that, clusterin has the potential to form amyloid-like fibrils. In fact, there have been reports of other proteins that function as molecular chaperones and exhibit amyloidogenicity *in vitro* (Ecroyd & Carver, 2009; Ecroyd *et al*., 2010) or *in vivo* (Niewold et al., 1999). Despite lacking any sequence similarity, this group of proteins shares a significant number of similar features, besides having the potential to form amyloid-like fibrils.

A prominent example is *α*-crystallin, a small heat shock protein (sHSP), which is highly expressed in the eye lens. Full-length *α*-crystallin has been found to form amyloid-like fibrils *in vitro* (Ecroyd & Carver, 2009) and is believed to be implicated in the formation of cataract (Meehan *et al*, 2004). As most sHSPs, *α*-crystallin exhibits remarkable structural plasticity, which is intertwined with its chaperone activity (Basha et al, 2012). Interestingly, *α*-crystallin’s chaperone activity is exhibited in a very similar manner to that of clusterin (Carver et al, 2003). sHSPs exist in solution as oligomeric structures, ranging from 9 to 50 subunits. Those structures disassociate under stress and release functional subunits (Haslbeck & Buchner, 2002). *α*-crystallin in particular, possesses a dynamic oligomeric structure, which, under increasing temperature, allows exchange of subunits within proximate oligomers. The so-called “traveling subunits” are believed to play a fundamental role in *α*-crystallin’s chaperone activity, by exposing hydrophobic surfaces and consequently, allowing hydrophobic interactions with misfolded proteins, including A*β* (Doss et al, 1997; Inoue et al, 2016; Narayan et al, 2012). Even though this is the physiological mechanism in which *α*-crystallin functions, it is reminiscent of how amyloidogenic proteins behave on their way to self-assembly and formation of amyloid fibrils (Cheon et al, 2007). This might explain the reports of *α*-crystallin forming amyloid-like fibrils *in vitro* (*Ecroyd & Carver, 2009*).

Another relevant example is that of the caseins, a group of proteins that comprise the largest part of the total protein content in milk (Kunz & Lonnerdal, 1990) and are usually found in aggregates, known as casein micelles. The micelles are primarily composed of four major proteins, namely *α*_s1_-, *α*_s2_-, *β*- and *κ*-casein (McMahon & Oommen, 2008). All four of them have been classified as intrinsically disordered proteins, because of the considerable structural flexibility that they exhibit (Redwan et al, 2015). In consistence with clusterin and *α*-crystallin, the major casein proteins are believed to function as molecular chaperones (Morgan *et al*, 2005), while *κ*-casein forms amyloid fibrils *in vitro* (Ecroyd et al., 2010) and *α*_s2_-casein forms amyloid fibrils *in vivo*, in corpora amylacea, in the bovine mammary gland (Benson et al., 2020; Niewold et al., 1999). Yet, in physiological conditions, *α*_s1_-casein interacts with *α*_s2_-casein, forming *α*_s_-casein, effectively halting the formation of amyloid fibrils (Thorn et al, 2008). Similarly, *α*_s_-casein and *β*-casein inhibit *κ*-casein amyloid formation, by shielding monomeric *κ*-casein hydrophobic surfaces with similar regions of their respective sequences (Thorn *et al*, 2005).

In summary, clusterin, *α*-crystallin, and the casein proteins share common features:

1. They function as molecular chaperones, by exposing hydrophobic regions, interacting with and stabilizing misfolded proteins on their way to aggregation.
2. They show structural plasticity, either having molten globule-like or intrinsically disordered regions on their sequence.
3. They are found in solution as heterogeneous aggregates, oligomers or micelles, which dissociate under conditions of stress.
4. They all interact with one, or more amyloidogenic proteins.
5. Parts of their sequence, or the full-length proteins, are known to form amyloid-like fibrils *in vitro* or *in vivo*.

These similarities may underlie a common mechanism in which the cell prevents toxic amyloid formation under different conditions of stress, by utilizing the intrinsic amyloidogenicity of specific molecular chaperones. For clusterin, amyloidogenesis inhibition could be activated under pH-induced stress and achieved through the interaction of its aggregation-prone regions with amyloidogenic proteins, on their way to aggregation. Likewise, *α*-crystallin could act under temperature-induced stress and the casein proteins may present a more specific system of amyloid inhibition, present primarily at the mammary gland.

### A putative mechanism for clusterin-mediated inhibition of *Aβ* amyloidogenesis

Clusterin *α*-chain peptide-analogues can inhibit A*β* amyloidogenesis *in vitro*, and the structural flexibility that natively disordered regions introduce, may enable them to act semi-independently of the protein. Having established that, we opted to propose a mechanism in which clusterin could use its aggregation-prone regions to exert its molecular chaperone activity on A*β*.

It is believed that A*β* amyloidogenesis is promoted by acidic pH (Su & Chang, 2001). As previously mentioned, mildly acidic pH also allows the disassociation of clusterin’s heterogeneous aggregates, releasing chaperone-active subunits (Poon et al., 2002). In its disassociated form, clusterin is assumed to hydrophobically interact with its client-proteins through amphipathic *α*-helices, which are surrounded by molten globule-like regions (*Bailey et* al., 2001). ^12^NFHAMFQ^18^, one of clusterin’s aggregation-prone regions, is located on a putative amphipathic *α*-helix. However, we have already proved that NFHAMFQ also has the propensity to form *β*-strands (Tsiolaki et al., 2017), hinting that the broader region has the intrinsic conformational properties of a chameleon sequence. When in proximity with an A*β* fibril, the *α*-helical ^12^NFHAMFQ^18^ could unfold and then refold into a *β*-strand, allowing interaction with A*β*’s *β*-stranded regions and temporarily halting fibril growth. This process could trigger the exposition of clusterin’s other aggregation-prone regions and allow it to attach itself at the edge of the A*β* fibril, via *β*-strand hydrogen bonding. In this manner, it would not allow A*β* monomers to elongate the fibril, effectively blocking further polymerization. At the same time, free clusterin subunits could bind to A*β* monomers, forming complexes and stabilizing their structure, until it passes the baton to other molecular chaperones, once again, halting fiber growth and minimizing the possibility of toxic secondary nucleation (Beeg et al, 2016). The suggested mechanism is visualized in Fig 7.

**Figure 7.**
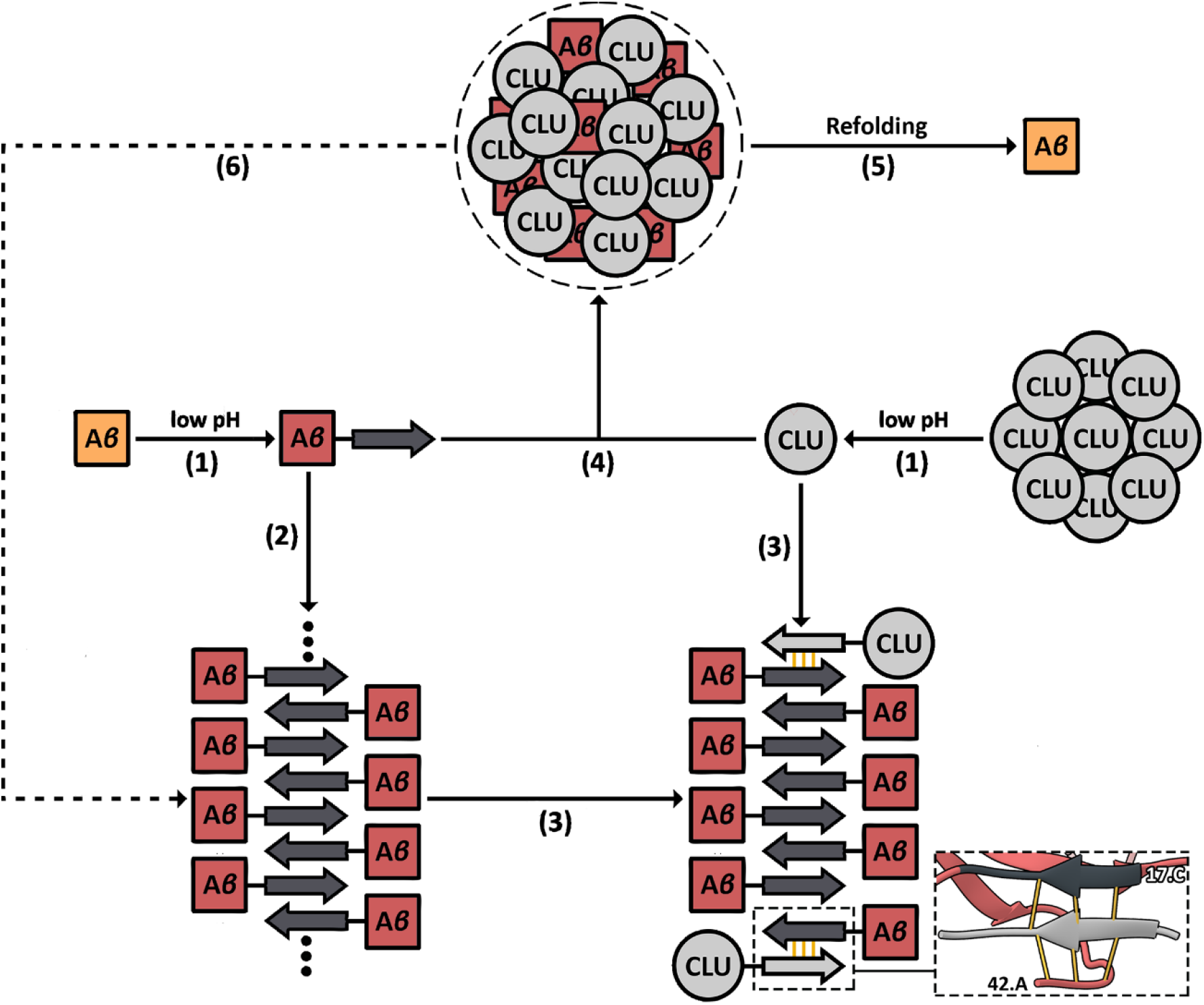
Putative mechanism of clusterin-mediated inhibition of A*β* amyloidogenesis. (1) Under acidic pH-induced stress, A*β* may partially unfold, exposing hydrophobic regions with the propensity to form *β*-strands. At the same time, the acidic pH triggers the disassociation of clusterin’s heterogeneous aggregates. (2) Misfolded A*β* begin to form amyloid fibrils, but (3) the disassociated clusterin subunits (CLU) halt fibril growth, utilizing their aggregation-prone regions, which tend to form *β*-strands as well. The outlined picture is derived from the last frame of the molecular dynamics simulation of A*β*_42_+EVVVKLF. (4) Clusterin binds misfolded A*β* monomers, forming complexes and stabilizes their structure, until (5) they are refolded by other molecular chaperones. (6) In case misfolded A*β* species are produced faster than clusterin can stabilize their structure, it essentially brings them into proximity with remote monomers, accelerating amyloid formation.

The putative mechanism that was just described, is also supported by the identification of a pathological mutation on clusterin’s gene, in AD patients. As previously mentioned, p.I360N changes the aggregation-prone region ^81^ILSVD^85^ to ^81^NLSVD^85^ (Bettens et al., 2015). Considering that out of the peptides we examined, ILSVD is hinted to be one of the better A*β* fibril formation inhibitors, this mutation could lead to a malfunction in the clusterin-mediated amyloidogenesis inhibition system. Thus, the pathogenicity of the phenotype could be explained by clusterin’s inability to inhibit amyloid formation, due to the absence of its most aggregation-prone region, that being ^81^ILSVD^85^, which would normally inhibit A*β* fibril formation.

The aforementioned mechanism can explain why clusterin is found co-localized with A*β* fibrillar deposits (Calero et al., 2005) and is consistent with reports of clusterin decelerating A*β* fibril formation (Beeg et al., 2016; de Retana et al, 2019). However, clusterin’s occasional contribution to the appearance of AD symptoms (DeMattos et al., 2002; Oh et al., 2019) needs further elucidation. It is logical to assume that, when A*β* misfolded monomers are produced faster than what clusterin’s existing population can process, the resulting complexes would fail to stabilize A*β*’s structure, essentially bringing remote misfolded monomers into proximity. This could accelerate amyloid fibril formation (Fig 6 – 6), while clusterin would end up co-localized with the fibrillar deposits. This hypothesis is supported by the findings of Yerbury *et al*., which show that clusterin’s enhancing effect on amyloid formation is caused by low clusterin:substrate ratio (Yerbury et al., 2007). A relevant example could be found in patients with familial Alzheimer’s disease (FAD), where A*β* is produced much faster than usual (Wu et al, 2012). In fact, most pre-clinical studies on AD use FAD animals as models, because it is a guaranteed method of acquiring early-onset AD test subjects (LaFerla & Green, 2012). This could also be the case for Oh *et. al*, who showed that clusterin-null 5XFAD mice, show AD symptoms later than their littermate counterparts (Oh et al., 2019). In the end, molecular chaperone-mediated amyloidogenesis inhibition, seems to be a double-edged sword for the cell, effective, but at the same time, conditionally harmful. The inadvertent bias that pre-clinical studies introduce, using test subjects with FAD, rather than sporadic AD, could make the inhibition system seem much more flawed than it really is.

### Peptide-based amyloidogenesis inhibitors, derived from aggregation-prone protein regions

We ultimately proposed that the cell can prevent amyloid formation by utilizing the intrinsic amyloidogenicity of specific molecular chaperones, similar to how functional amyloids work. If that statement proves to be true, harnessing the ability of molecular chaperones to halt amyloid formation could be essential to tackling AD and other amyloidoses. The key to inhibiting amyloid formation lies in the aggregation-prone regions of said proteins, thus synthesizing and studying peptides from such regions could provide novel inhibitors of A*β* amyloidogenesis and amyloid formation in general (Connelly et al, 2010).

It is worth mentioning that, the ability of molecular chaperones to inhibit amyloid formation doesn’t seem to be substrate-specific, meaning that one chaperone could act as an amyloidogenesis inhibitor for a variety of different amyloidogenic proteins, even if they are not known to interact *in vivo*. For example, though *α*-crystallin is known to interact with A*β* (Narayan et al., 2012), the casein proteins are not. Despite that, all of them can inhibit A*β* amyloidogenesis *in vitro* (*Carrotta et al, 2012; Raman et al, 2005*). Furthermore, this also seems to be the case for aggregation-prone peptides that are not derived from regions of molecular chaperones (Lu et al., 2019). Mimicking the physiological system for amyloidogenesis prevention and extending it to aggregation-prone peptide-analogues of non-chaperone-active proteins could provide us with a much greater variety of potential inhibitors.

Despite the potential that specific peptides show as amyloidogenesis inhibitors, it is of vital importance to ensure that they are non-toxic for the cell. It is noteworthy that, even though some molecular chaperones have amyloidogenic properties, they seem to not have negative effects on the survival of the cell. In a similar manner, functional amyloids are not harmful to the organisms in which they appear. It is believed that controlling the expression of amyloidogenic proteins, regulating fibril formation with co-localized proteins-molecules and the fact that they are often found isolated inside membrane bound compartments, are some of the reasons why functional amyloids are not toxic (Jackson & Hewitt, 2017). For the peptide-inhibitors to be usable, these questions must be accurately answered, and the findings should be incorporated in their implementation as potential drugs for amyloidoses, especially when it comes to peptide concentration and the complementary substances that should be administered.

## Materials and Methods

### Prediction of potential aggregation-prone and natively disordered regions in human clusterin

AMYLPRED (Frousios et al., 2009), a consensus algorithm for the prediction of amyloid propensity, was used for the identification of aggregation-prone regions in wild-type, mature clusterin (Uniprot AC: P10909). The *α*-chain, which is our focus in the current study, has five such regions. The first one (^12^NFHAMFQ^18^) has previously been studied by our lab and proved to exhibit amyloidogenic properties *in vitro* (Tsiolaki et al., 2017). This time, we opted to study the remaining four, those being ^31^MDIHF^35, 82^LSV^84, 155^YYLRVTT^161^ and ^177^EVVVKLF^183^. Natively disordered regions were predicted using PONDR^®^ VLTX (http://www.pondr.com).

### Peptide synthesis

All peptide-analogues were synthesized and lyophilized by GeneCust^©^ Europe, France. The exact sequences for the peptide-analogues were NFHAMFQ, AMDIHF, ILSVD, NLSVD, YYLRVTT and EVVVKLF. The regions that were used for the second and third peptide-analogues were ^30^AMDIHF^35^ and ^81^ILSVD^85^, which are extensions of the original predicted regions. NLSVD is a mutant counterpart of ILSVD. The purity of all the synthesized peptides was higher than 98%, with the N- and C-terminals being free.

A*β*_42_ was also synthesized and lyophilized by GeneCust^©^ Europe, France. The exact sequence is DAEFRHDSGYEVHHQKLVFFAEDVGSNKGAIIGLMVGGVVIA. The purity of the peptide was higher than 95%, with the N- and C-terminals being free.

### Sample preparation

#### Disaggregation of pre-existing aggregates

Each lyophilized peptide was dissolved in 1,1,1,3,3,3-Hexafluoro-2-propanol (HFIP; Sigma^©^) at a concentration of 1 mg/mL. All clusterin peptides were mixed with equimolar amounts of A*β*_42_. Individual peptide solutions and A*β*_42-_ clusterin peptide solutions were left to dry overnight in a fume hood, at room temperature, until thin peptide-containing films were created. The peptide-containing films were stored at −20 °C.

#### *In vitro* fibril formation

The peptide-containing films were left at room temperature for 30 minutes and dissolved in 0.1 M HEPES buffer (pH=7.4; Sigma^©^) or dH_2_O (pH=5.57), at concentrations ranging from 10 μM to 1 mM. The peptide solutions were incubated for 1 week at 37 °C, unless stated otherwise.

### Transmission Electron Microscopy (TEM)

A 3 μL droplet of 100 or 500 μM peptide solution was applied to glow-discharged 400-mesh carbon-coated copper grids for 60 seconds. Directly afterwards, the grids were stained with a 3 μL droplet of 2% (w/v) aqueous uranyl acetate for another 60 seconds. Excess stain was blotted away with filter paper and air-dried. The grids were examined with a Morgagni^™^ 268 transmission electron microscope, operated at 80 kV. Digital acquisitions were performed with an 11 Mpixel side mounted Morada CCD camera (Soft Imaging System, Muenster, Germany).

### X-ray diffraction from oriented protein fibers

#### Oriented fiber formation

1 mM peptide solutions were incubated for two weeks, to form viscous solutions, which facilitate the formation of oriented fibers. A 10 μL droplet of each peptide solution was placed between two quartz capillaries covered with wax, spaced approximately 1.5 mm apart and mounted horizontally on a glass substrate, as collinearly as possible. The droplet was allowed to dry for 3 days, until oriented fibers were formed.

#### X-ray diffraction and analysis

The oriented fibers were shot with X-rays and diffracted. The X-ray diffraction pattern was collected using a SuperNova-Agilent Technologies X-ray generator, equipped with a 135 mm ATLAS CCD detector and a 4-circle kappa goniometer, at the Institute of Biology, Medicinal Chemistry and Biotechnology, National Hellenic Research Foundation (CuKα high intensity X-ray micro-focus source, *λ* = 1.5418 Å), operated at 50 kV, 0.8 mA. The specimen-to-film distance was set at 52mm and the exposure time was set to 200s. The X-ray patterns were initially viewed using CrysAlisPro (CrysAlis^PRO^, 2014) and afterwards measured with iMosFLM (Battye et al, 2011).

### Attenuated total reflectance Fourier-transform infrared (ATR FT-IR) spectroscopy

A 10 μL droplet of 500 μM peptide solution was cast on a front-coated Au mirror and left to dry slowly at room temperature, until a thin peptide-containing film was created. Infrared spectra were obtained from these films at a resolution of 4 cm^-1^, utilizing an IR microscope (IRScope II by Bruker Optics) equipped with a Ge attenuated total reflectance (ATR) objective lens (20x) and attached to a Fourier-transform infrared (FT-IR) spectrometer (Equinox 55, by Bruker Optics). Ten 32-scan spectra were collected from each sample and averaged to improve the sound/noise (S/N) ratio. Both are shown in the absorption mode after correction for the wavelength dependence of the penetration depth (pd-*λ*). Absorption band maxima were determined from the minima in the second derivative of the corresponding spectra. Derivatives were computed analytically using routines of the Bruker OPUS/OS2 software, including smoothing over a ±13 cm^-1^ range around each data point, performed by the Savitzky–Golay (Savitzky & Golay, 1964) algorithm. Data was visualized using OriginPro 7 (OriginLab Corporation, Northampton, MA, USA).

### Congo Red Birefringence Assay

A 3 μL droplet of the 500 μM peptide solution was applied to glass slides and stained with a 10 mM Congo Red (Sigma^©^) solution in PBS (137 mM NaCl, 27 mM KCl, 100 mM Na_2_HPO_4_, 18 mM KH_2_PO_4_, pH=7.4) or dH_2_O for approximately 30 minutes. Then, they were washed several times with 90% ethanol and were left to dry approximately for 10 minutes. The samples were observed under bright field illumination and between crossed polars, using a Leica MZ7.5 polarizing stereomicroscope, equipped with a Sony *α*6000 camera.

### Thioflavin T (ThT) Kinetic Assay

ThT fluorescence measurements were conducted at 37 °C, in black 96-well plates with flat, clear bottoms, using a Tecan Spark microplate reader. The top of the plates was sealed with microplate covers and the fluorescence readings were performed through the bottom. A 444 nm filter was used for excitation and a 484 nm filter for emission. HFIP peptide films were dissolved in DMSO and diluted in HEPES for a final DMSO concentration of less than 5% v/v. The reaction solutions contained freshly prepared 10 μM disaggregated peptide solutions (individual A*β*_42_ or A*β*_42_-clusterin peptide solutions) and 25 μM ThT (Sigma^©^) in dH_2_O. ThT background fluorescence was measured in the absence of peptide solutions. Each experiment was conducted in triplicates. Measurement lasted for 50 hours and fluorescence readings were collected every 15 minutes. ThT background fluorescence was subtracted from the peptide fluorescence readings at each time point. Standard deviation was calculated, and the data was normalized. 100 arbitrary units correspond to maximum individual A*β*_42_ fluorescence intensity. Data was visualized using RStudio (package ggplot2).

### Molecular Dynamics Simulations

#### Structure acquisition and molecular docking

Simulations were performed using an NMR structure of pentameric A*β*_42_ (PDB ID: 2BEG) (Luhrs et al., 2005), while a structure for EVVVKLF was generated through the “Builder” tool in PyMOL (Delano, 2005). A*β*_42_-EVVVKLF docking was performed through the automated protein docking server ClusPro (Kozakov et al., 2013; Kozakov et al., 2017; Porter et al., 2017; Vajda et al., 2017). The generated clusters were evaluated with the balanced scoring scheme.

#### Molecular dynamics simulations

The derived complex was subjected to molecular dynamics simulations via GROMACS v. 2018.1 (Kutzner et al, 2019). The simulations employed the AMBER99SB-ILDN protein, nucleic AMBER94 force-field (Lindorff-Larsen et al, 2010). The complex was placed in a 1.2 nm cubic box of 3-point model (TIP3P) water (Jorgensen, 1983) and the system was ionized using NaCl molecules, mimicking neutral pH conditions. Each simulation system was subjected to energy minimization, in a maximum of 2000 steps, using the steepest descent algorithm, followed by two stages of equilibration simulations with position restraints applied on protein coordinates. Specifically, a 100 ps simulation was performed in the canonical (NVT) ensemble to equilibrate temperature at 310 K, using the Berendsen-thermostat (Berendsen, 1984). Following the first equilibration, a 100 ps simulation was performed in the isothermal-isobaric (NPT) ensemble to control pressure isotopically at 1.013 bar (1 atm), using the Berendsen weak coupling algorithm (Berendsen, 1991) and the Berendsen-thermostat at 310 K. Finally, the MD simulation with position restraints removed was carried out for 600ns at 310K. One control run of individual A*β*_42_ was also performed. Periodic boundary conditions were applied to all directions. The LINCS algorithm (Berk Hess, 1997) was applied to model bond constraints, enabling the use of a 2 fs time-step. Short range non-bonded interactions were modeled using a twin-range cutoff at 0.8 nm, while long-range electrostatic interactions were modeled using the Particle Mesh Ewald (PME) method, with a Fourier grid spacing at 0.12 nm (Essmann, 1995).

#### Analysis of simulation results

Simulation results were analyzed using various GROMACS utilities, and Visual Molecular Dynamics (VMD) v. 1.9.4 (Humphrey et al, 1996). Structural stability was checked using the “rms” tool (Maiorov & Crippen, 1995). Secondary structure was calculated using the “do_dssp” tool (Kabsch & Sander, 1983; Touw et al., 2015). Hydrogen bonds were calculated using the “hbond” tool (van der Spoel et al, 2006). Frames were extracted every 100ps and used for the analysis. Pictures were collected with UCSF Chimera (Pettersen et al., 2004) or PyMOL (Delano, 2005).

## Availability of published material and data

This study includes no data deposited in external repositories.

## Acknowledgments

This research has been co-financed by the European Union and Greek national funds through the Operational Program “Competitiveness, Entrepreneurship and Innovation”, under the call “RESEARCH – CREATE – INNOVATE” (project code: T1EDK-00353). We acknowledge support of this work by the project “INSPIRED-The National Research Infrastructures on Integrated Structural Biology, Drug Screening Efforts and Drug target functional characterization” (MIS 5002550) which is implemented under the Action “Reinforcement of the Research and Innovation Infrastructure”, funded by the Operational Program “Competitiveness, Entrepreneurship and Innovation” (NSRF 2014-2020) and co-funded by Greece and the European Union (European Regional Development Fund). The molecular dynamics simulations were performed utilizing computational time granted from the Greek Research & Technology Network (GRNET) at the National HPC facility – ARIS under project ID “PR007003-AbetaDynamics”. We also acknowledge Dr. George Baltatzis for kindly assisting us with handling the Morgagni Microscope at the 1st Department of Pathology, Medical School, National and Kapodistrian University of Athens. Finally, we acknowledge the support of Dr. Aimilia Sklirou, who kindly assisted with handling the Tecan Spark microplate reader.

## Author contributions

P.M.S. performed sample preparation, Congo Red birefringence and ThT kinetics assays, acquired and processed TEM, X-ray diffraction and ATR FT-IR spectroscopy data, analyzed the molecular dynamics simulations results, helped in conceiving the project and wrote the initial draft of the manuscript. G.I.N. participated and assisted in all the experiments. P.L.T. co-supervised and helped in conceiving the project. M.K.T. performed the molecular dynamics simulations and participated in the analysis of the results. N.C.P. supervised the molecular dynamics simulations. I.P.T. helped in conceiving the project and supervised the ThT kinetics assay. V.A.I. supervised and helped in conceiving the project. All authors discussed the results and edited or commented on the manuscript.

## Expanded View

**Figure EV1.**
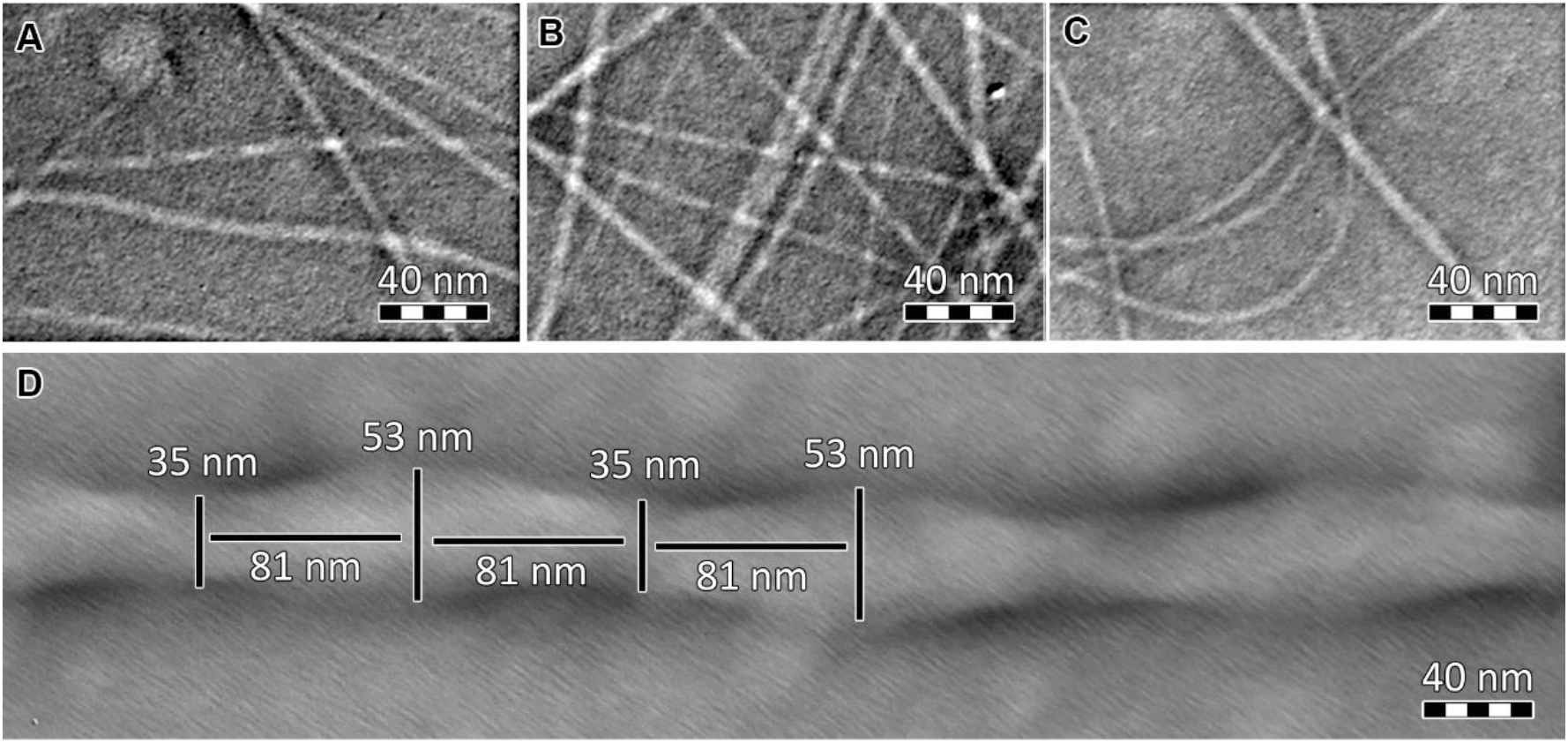
Electron micrographs of polymorphic structures formed by AMDIHF and EVVVKLF amyloid-like fibrils, after 7 days of incubation, at 500 μM. (A) Supercoil formed by AMDIHF amyloid-like fibrils. (B) Tape formed by laterally interacting AMDIHF amyloid-like fibrils. (C) An amyloid-like fibril formed by AMDIHF, with a diameter of approximately 160 Å, is split into two 80 Å fibrils. (D) EVVVKLF amyloid-like fibrils form large, twisting ribbons, with a 162 nm pitch, a minimum width of 35 nm, and a maximum width of 53 nm.

**Figure EV2.**
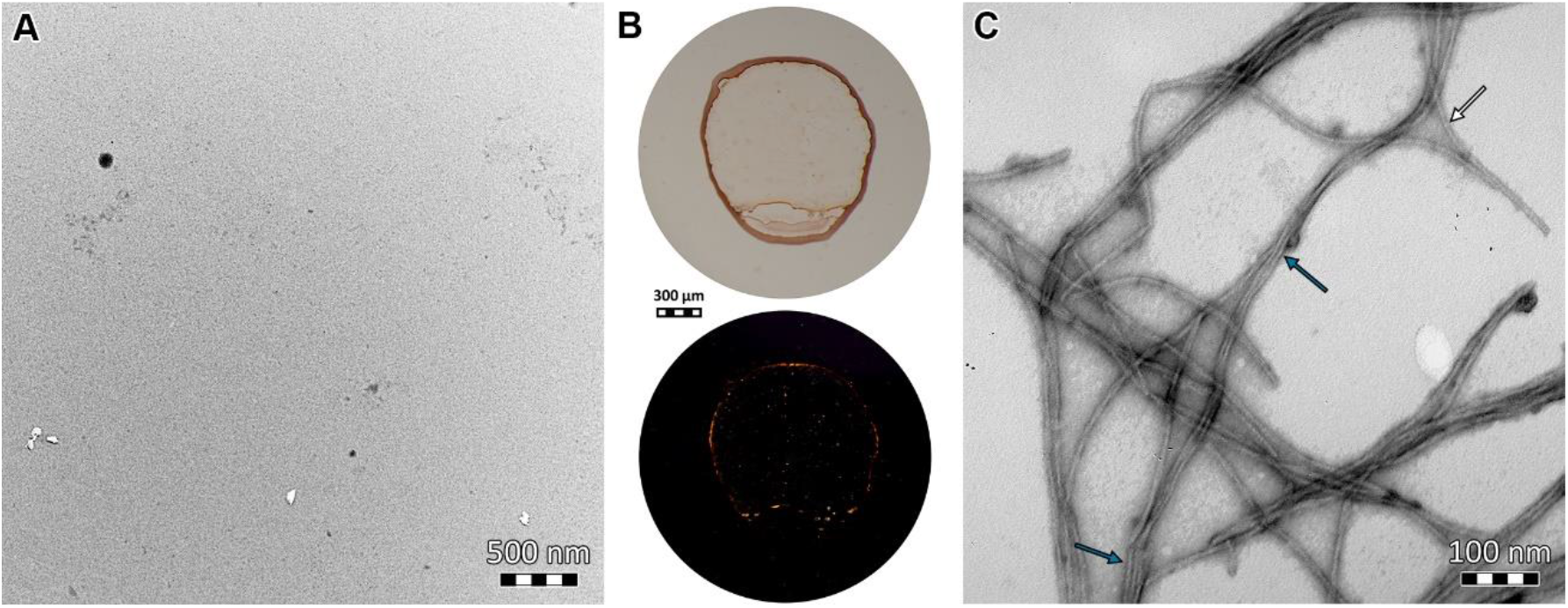
The mutant NLSVD does not form amyloid-like fibrils and does not inhibit A*β*_42_ fibril formation. (A) Electron micrographs reveal that NLSVD 500 μM solution does not contain amyloid-like fibrils, after 7 days of incubation. (B) Photomicrographs of gels derived from NLSVD 500 μM solution. Congo red is faintly bound, as seen under bright field illumination (upper) and apple-green birefringence is not exhibited under crossed polars (lower). (C) Electron micrographs of A*β*_42_+NLSVD (100 μM A*β*_42_ and 100 μM NLSVD) reveal that NLSVD does not inhibit amyloid fibril formation, after 7 days of incubation. A*β*_42_ fibrils (white arrow) are wider than those of individually incubated A*β*_42_ fibrils (Figure 4d) and have a higher tendency to interact laterally and form loosely bound tapes (blue arrows).

